# A quaternion model for single cell transcriptomics

**DOI:** 10.1101/2022.07.21.501020

**Authors:** H. Robert Frost

**Affiliations:** Department of Biomedical Data Science, Geisel School of Medicine, Dartmouth College, Hanover, NH 03755, USA

## Abstract

We present an approach for modeling single cell RNA-sequencing (scRNA-seq) and spatial transcriptomics (ST) data using quaternions. Quaternions are four dimensional hypercomplex numbers that, along with real numbers, complex numbers and octonions, represent one of the four normed division algebras. Quaternions have been primarily employed to represent three-dimensional rotations in computer graphics with most biomedical applications focused on problems involving the structure and orientation of biomolecules, e.g., protein folding, chromatin conformation, etc. In this paper, we detail an approach for mapping the cells/locations in a scRNA-seq/ST data set to quaternions. According to this model, the quaternion associated with each cell/location represents a vector in ℝ^3^ with vector length capturing sequencing depth and vector direction capturing the relative expression profile. Assuming that biologically interesting features of an scRNA-seq/ST data set are preserved within a rank three reconstruction of the unnormalized counts, this representation has several benefits for data analysis. First, it supports a novel approach for scRNA-seq/ST data visualization that captures cell state uncertainty. Second, the model implies that transformations between cell states can be viewed as three-dimensional rotations, which have a corresponding representation as rotation quaternions. The fact that these rotation quaternions can be interpreted as cells enables a novel approach for characterizing cell state transitions with specific relevance to the analysis of pseudo-temporal ordering trajectories. Most importantly, a quaternion representation supports the genome-wide spectral analysis of scRNA-seq/ST data relative to a single variable, e.g., pseudo-time, or two variables, e.g., spatial coordinates, using a one or two-dimensional hypercomplex discrete Fourier transform. An R package supporting this model and the hypercomplex Fourier analysis of ST data along with several example vignettes is available at https://hrfrost.host.dartmouth.edu/QSC.

## 1 Background

### 1.1 Quaternions

Quaternions are four dimensional hypercomplex numbers that, along with real numbers, complex numbers and octonions, represent one of the four normed division algebras. Formally, hypercomplex numbers are elements of a finite-dimensional algebra over the real numbers that is unital but may not be associative or commutative [1] and have a general representation given by:

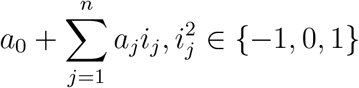

where *a*_0_ and the *a*_*j*_ are real numbers and the *i*_*j*_ form an n-dimensional basis. Following this representation, *n* = 0 for real numbers, *n* = 1 for complex numbers, *n* = 3 for quaternions and *n* = 7 for octonions with 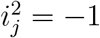, i.e., real numbers, complex numbers, quaternions and octonions are the hypercomplex numbers with dimensions 1, 2, 4 and 8 that are each comprised by a real number and a vector in an n-dimensional imaginary space. While real and complex numbers are commonly employed in applied mathematics to model experimental data, quaternions, and especially ontonions, are infrequently used outside of a theoretical context.

Quaternions were first explored by Gauss in 1809, however, because that work was not published until 1900, the discovery of quaternions is usually credited to the Irish mathematician William Hamilton in 1843 [2]. Hamilton’s discovery was motivated by attempts to extend complex numbers to three dimensions. In his work on three dimensional hypercomplex numbers, Hamilton struggled with multiplication/division, i.e., what is the quotient of two points in ℝ^3^? Hamilton realized these problems could be solved by moving to four dimensional hypercomplex numbers and, when he had this realization, famously carved the formula for the quaternion imaginary basis vectors into the Brougham Bridge:

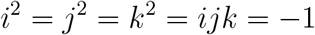

More specifically, a quaternion *q* is a four dimensional hypercomplex number defined by (1).

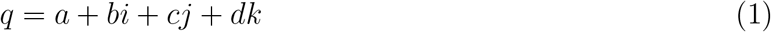

where {*a, b, c, d*} ∈ ℝ and *i, j*, and *k* form a three dimensional imaginary basis with:

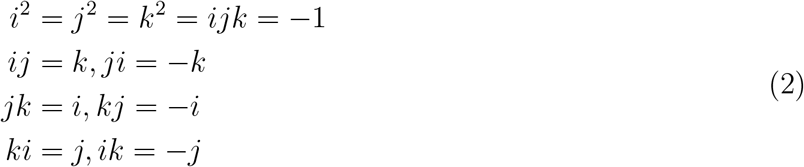

Addition and subtraction of quaternions is simply performed by adding and subtracting the associated terms. For example, if *q*_1_ = *a*_1_ + *b*_1_*i* + *c*_1_*j* + *d*_1_*k* and *q*_2_ = *a*_2_ + *b*_2_*i* + *c*_2_*j* + *d*_2_*k*, then *q*_1_ + *q*_2_ = *a*_1_ + *a*_2_ + (*b*_1_ + *b*_2_)*i* + (*c*_1_ + *c*_2_)*j* + (*d*_1_ + *d*_2_)*k*. Multiplication of quaternions follows the distributative law and the rules for multiplication of the basis vectors given by (2). An important consequence of (2) is that multiplication of quaternions is not commutative, e.g., *ij* ≠ *ji*. Division of quaternions is defined by the pre or post multiplication by the inverse of the denominator, which, given the noncommunicativity of multiplication, may be distinct. So, *q*_1_*/q*_2_ has two possible values: 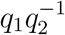 or 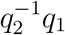. The inverse of a quaternion is defined as the ratio of the conjugate (*q*\*) and the square of the norm (||*q*||):

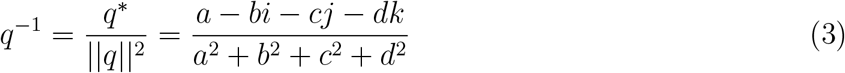

It is often convienent to separate quaternions into scalar (*r* = *a*) and vector 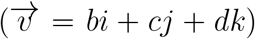 parts:

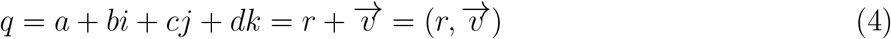

If *b* = *c* = *d* = 0, the quaternion is referred to as a scalar quaternion. If *a* = 0, the quaternion is referred to as a vector or pure quaternion. Using the scalar/vector notation, multiplication of quaternions can be conveniently represented using vector dot and cross product notation:

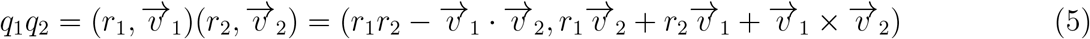

Interpretation of (5) can be aided by considering two special cases:

1. Multiplication by a scalar: If *q*_2_ is a scalar quaternion, i.e., 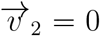, then the multiplication simplifies to the scaling of *q*_1_ by the scalar part of *q*_2_:

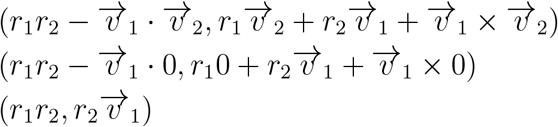
2. Multiplication of two vector quaternions: If both *q*_1_ and *q*_2_ are vector quaternions, i.e., *r*_1_ = *r*_2_ = 0, then multiplication simplifies to just the dot and cross product terms:

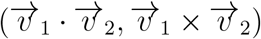

If 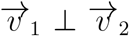, then 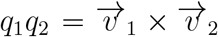, which is a vector quaternion that is orthogonal to the plain defined by *q*_1_ and *q*_2_. If 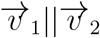, then 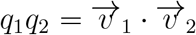, which is a scalar quaternion.

Following their introduction by Hamilton, quaternions were widely used in geometry and physics until the development of vector analysis methods. While interest in quaternions is now primarily restricted to pure mathematics, they are still employed for the efficient computation of 3D rotations, as detailed in Section 1.2 below.

### 1.2 Quaternion representation of three-dimensional rotations

An important feature of quaternions is their ability to efficiently represent rotations in ℝ^3^ [3]. According to Euler’s rotation theorem, any single rotation (or sequence of rotations) in ℝ^3^ can be modeled as a rotation of angle *θ* about an axis (the Euler axis) that can be represented by a unit vector. Let the ℝ^3^ vector 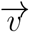 represent a vector from the origin to the point (*v*_*x*_, *v*_*y*_, *v*_*z*_) and model 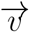 using the vector quaternion *q*_*v*_ = *v*_*x*_*i* + *v*_*y*_*j* + *v*_*z*_*k*. The rotation of 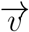 by angle *θ* about the axis defined by unit vector 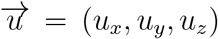 can be represented by the rotation quaternion *r* = *cos*(*θ/*2) + *sin*(*θ/*2)(*u*_*x*_*i* + *u*_*y*_*j* + *u*_*z*_*k*) and computed mathematically as:

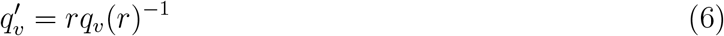

The coeffients of the generated vector quaternion 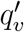 capture the coordinates of the rotated vector 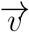 in ℝ^3^. The product of multiple rotation quaternions generates a quaternion whose orientation is produced by the corresponding sequence of rotations. There are several important benefits of the quaternion model for computing rotations relative to the vector analysis approach that employs a 3 × 3 rotation matrix:

- Rotation quaternions provide a more compact representation (4 numbers vs 9 for a rotation matrix)
- Rotation quaternions are a more interpretable representation (i.e., easy to translate between desired rotation and rotation quaternion)
- Rotation quaternions support efficient calculation of smooth rotations by robustly dividing a single large rotation into a sequence of small rotations.

These benefits have led to the frequent use of quaternions in computer graphics.

### 1.3 Analysis of biomedical data using quaternions

Given the advantages of quaternions for modeling rotations, most biological applications of quaternions have involved the structural analysis of biomolecules including problems involving protein folding [4], chromatin structure [5], and molecular dynamics [6]. Outside of these structural analysis problems, use of quaternions for the analysis of biological data has been very limited with the only notable developments involving the discovery of repetitive elements within either one-dimensional sequences (e.g., DNA) or two-dimensional arrays (e.g., images). For example, Brodzik explored the use of quaternions for the detection of tandem repeats in DNA using a periodicity transformation on a novel mapping of DNA bases to quaternions [7], and, in the space of image analysis, Ell and Sangwine [8] explored the mapping of RGB pixel values into quaternions to support the analysis of color images using a two-dimensional hypercomplex Fourier transform.

### 1.4 Single cell transcriptomics

Although multicellular tissues are comprised by a complex set of cell types and states, high-throughput genome-wide profiling has historically been limited to bulk tissue assays. The genomic values captured by these techniques reflect the average across all cell types and states present in the sample and, when significant heterogeneity exists in the tissue sample, provide only a rough approximation of the true biological state. To address the limitations of bulk assays, researchers have recently developed a range of methods that support the single cell measurement of RNA expression (e.g., single cell RNA sequencing or scRNA-seq [9]), DNA sequence (e.g., single nucleus exome sequencing [10]), chromatin accessibility (e.g., single cell ATAC sequencing (scATAC-seq) [11]), epigenetic modifications (e.g., single cell bisulfite sequencing [12]) and protein abundance (e.g., Cellular Indexing of Transcriptomes and Epitopes by Sequencing (CITE-seq) [13]. Among these techniques, scRNA-seq has seen the most rapid development with state-of-the-art systems (e.g., 10x Genomics Chromium System [14]) now able to cost-effectively profile tens-of-thousands to hundreds-of-thousands of cells isolated from a single tissue sample. More recent technical advances have enabled the joint profiling of multiple omics modalities for dissociated cells (e.g., joint scRNA-seq/scATAC-seq and joint scRNA-seq/CITE-seq) and the spatial transcriptomics (ST) analysis of whole tissue samples at the resolution of single cells (e.g., MERFISH [15]) or small cell groups (e.g., 10x Genomics Visium technology [16]). Important applications of single cell transcriptomics include cataloging the cell types in a tissue [17, 18], discovering novel cell subtypes [19], reconstructing dynamic processes via pseudo-temporal ordering [20], and analyzing cell-cell signaling [21].

Although scRNA-seq and ST data has the potential to more accurately characterize the biological state of complex tissues, statistical analysis of this data is very challenging. Single cell transcriptomics is high-dimensional (transcript abundance is typically measured on 10, 000 or more genes), extremely sparse (90% or more gene counts are typically 0), and noisy (due to significant ampliplification bias and other technical factors). Single cell assays typically output a *n* × *p* matrix of transcript counts for *n* cells/locations and *p* protein-coding genes. The use of unique molecular identifiers (UMIs) means that these counts correspond to the number of unique mRNA molecules whose sequence aligns to a specific gene. Statistically, scRNA-seq/ST data is typically modeled using a Poisson or negative binomial distribution, which is sometimes extended to support zero-inflation. To enable the comparison of cells/locations with different library sizes (i.e., the total number of UMIs captured for a given cell/location), the raw scRNA-seq/ST count data is usually normalized to relative abundance values. One simply normalization approach (refered to as “log-normalization” in the Seurat framework [22]) that adjusts for library size takes the log of a pseudo count of 1 plus the ratio of the scaled counts for a specific gene to the total number of counts for the cell/location or 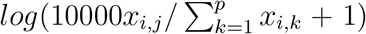, where *x*_*i,j*_ holds the UMI counts for gene *j* in cell/location *i*. More complex normalization methods that adjust for library size include the SCTransform method [23], which uses the residuals from a regularized negative binomial regression model of the raw counts on library size. Following normalization, some form of dimensionality reduction is typically employed (e.g., principal component analysis (PCA), uniform manifold approximation and projection (UMAP) [24], etc.) to make subsequent statistical analyses more tractable. Visualization of scRNA-seq/ST data is usually done using a projection onto the first two reduced dimensions.

## 2 Quaternion model for scRNA-seq/ST data

Our approach for mapping single cell transcriptomics (scRNA-seq or ST) data to quaternions is detailed in Algorithm 1 below. Although the focus on this paper is single cell transcriptomics, the same approach can be applied to other forms of count-based data, e.g., bulk RNA-seq.

### Algorithm 1 Quaternion model for scRNA-seq data

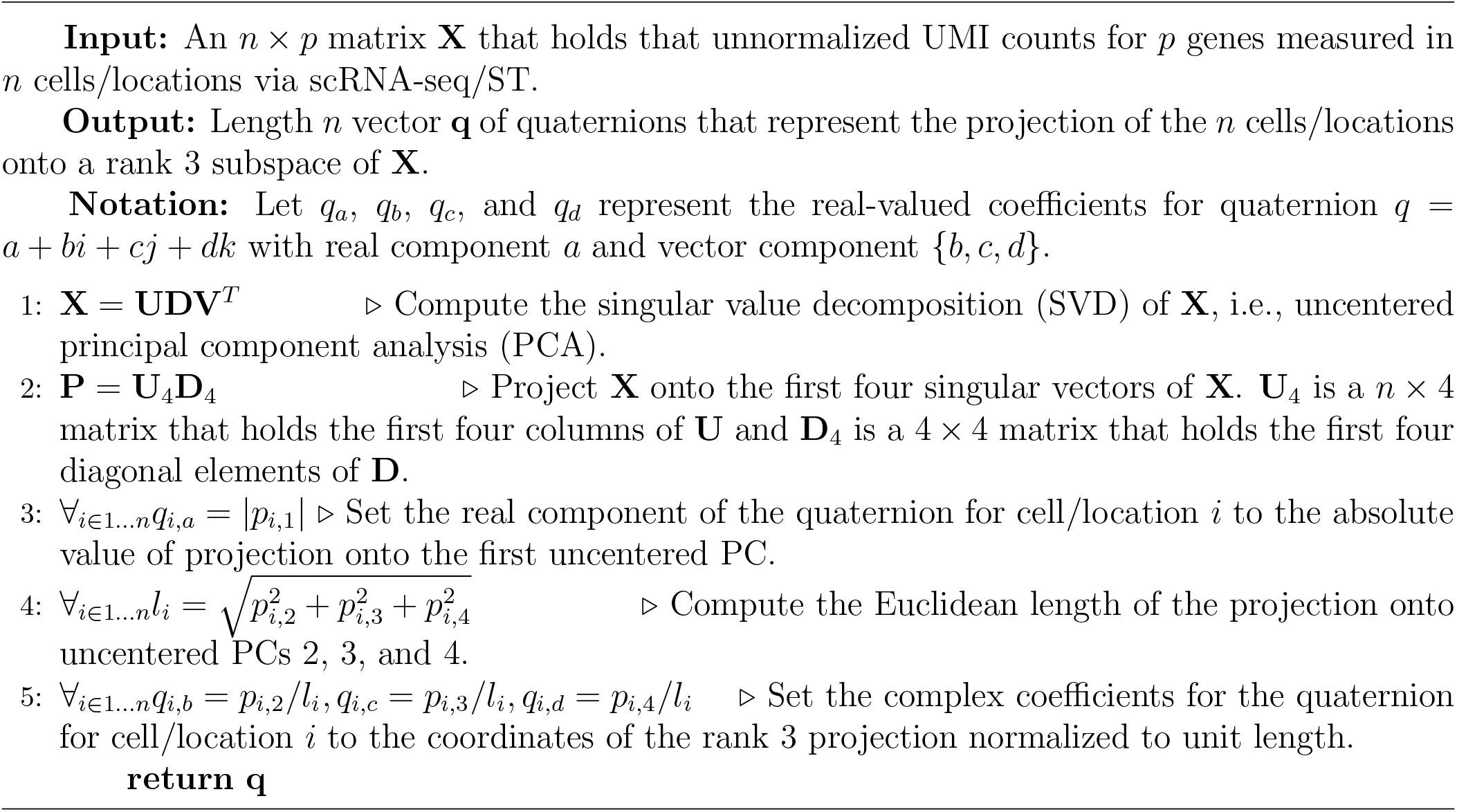

To improve the computational performance of Algorithm 1 on large scRNA-seq data sets, step 1 can be replaced by a randomized SVD algorithm (e.g., the randomized SVD implementation in the *rsvd* R package [25]) with only the first four singular vectors computed. The transformation of scRNA-seq and ST data into quaternions according to this model is supported by the QSC R package available at https://hrfrost.host.dartmouth.edu/QSC.

### 2.1 Properties of quaternion model

The quaternion scRNA-seq model outlined in Algorithm 1 projects the unnormalized count data for each cell onto the first four uncentered principal components (PCs). For unnormalized count data, the absolute value of the projection onto the first uncentered PC corresponds to library size, i.e., the sum of all counts for the cell, and the direction of the vector representing the projection of each cell onto the other uncentered PCs captures the cell’s relative expression profile. This model has the useful property that all cells with identical relative gene expression profiles fall onto a line that intersects the origin with distance along the line representing library size. Figure 1 below illustrates this phenomenon for simulated scRNA-seq relative abundance data for 5 genes and 30 cells split into three different populations. The first 10 cells were simulated with relative abundance values of {0.5, 0.2, 0.1, 0.1, 0.1}, the next 10 with relative abundance values of {0.1, 0.1, 0.1, 0.2, 0.5}, and the last 10 with relative abundance values of {0.1, 0.3, 0.3, 0.2, 0.1}. Within each population, the relative library size varied from 1 to 10. When these cells are projected onto the uncentered principal components, all cells sharing the same relative expression profile lie along a single line with distance from the origin proportional to library size.

**Figure 1:**
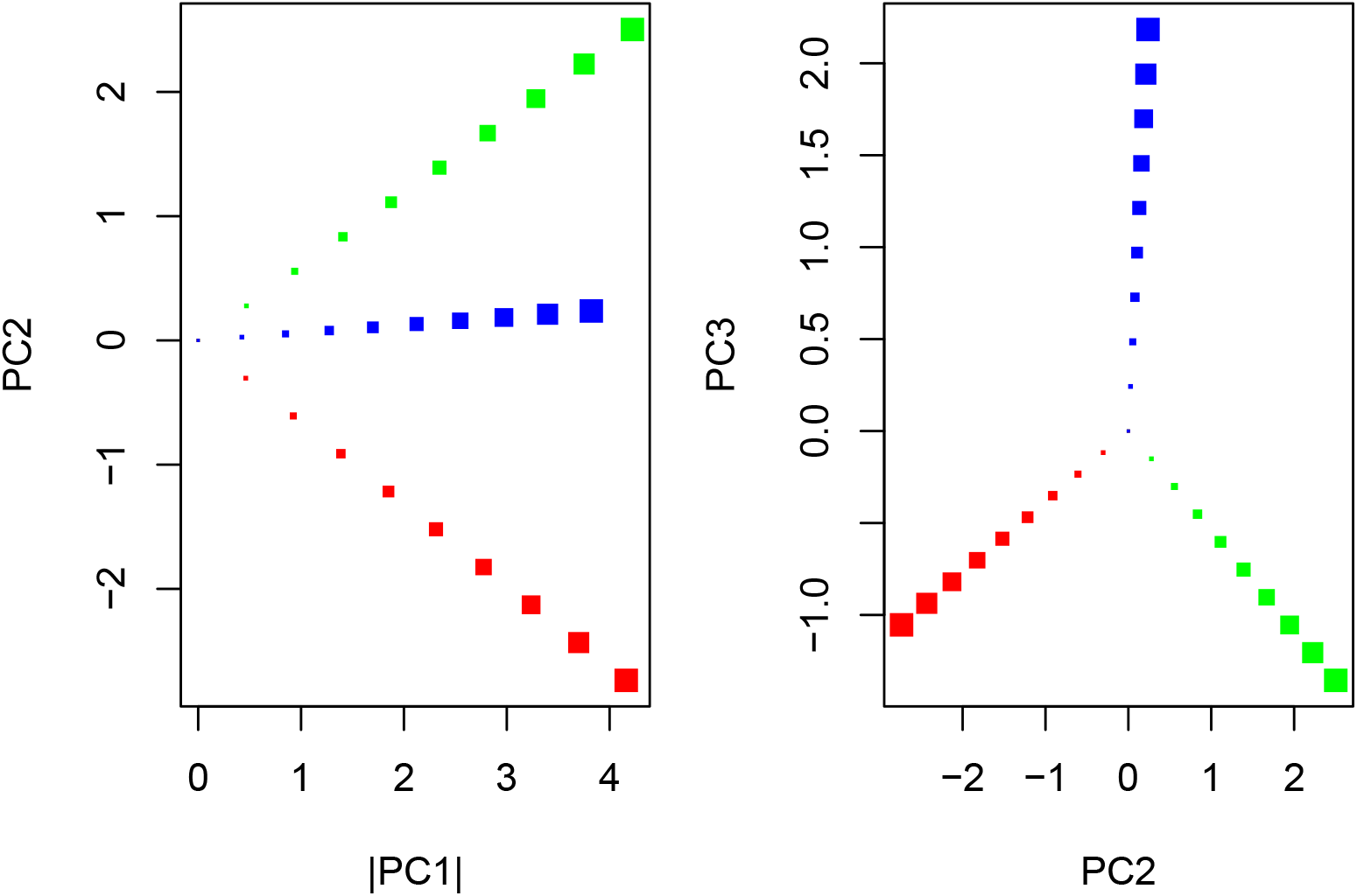
Projection of simulated scRNA-seq relative abundance data onto first three uncentered principal components. Points with the same color represent cells with the same relative abundance profile. Point size reflects the sum of all non-relative counts for the simulated cell.

Given these properties, the real part of the quaternion associated with each cell (as defined by Algorithm 1) represents library size and the vector part of the quaternion, which is normalized to unit length, represents the relative expression profile. Although this model only captures a rank two subspace of normalized scRNA-seq data, it captures the same data characteristics that are represented by standard two-dimensional (2D) visualizations of scRNA-seq data. The proposed quaternion model also has the useful property that transitions between different relative expression profiles are represented by a rotation in ℝ^3^, which is equivalent to the use of cosine similarity to capture the distance. Cosine similarity works well as a distance measure when the focus is on relative vs. absolute values (i.e., angle will be 0 for scaled versions of the same vector so is equivalent to comparing relative data) and is therefore appropriate for scRNA-seq and ST data that is typically normalized by library size to generate relative values.

### 2.2 Alternative quaternion models

The model detailed in Algorith 1 is only one potential mapping of scRNA-seq/ST data to quaternions and many other mappings are possible with the key limitation that a maximum of four independent dimensions can be represented. If capturing a rank four subspace of the normalized data is desired, the four quaternion coefficients could instead be set to the projection of each cell/location onto the first four centered PCs (or alternative dimensionality reduction technique, e.g., UMAP [24]) of the normalized data. While this form of mapping would capture a rank four subspace of the normalized data (as opposed to the rank two subspace captured by Algorithm 1), it lacks the clear interpretation of our proposed mapping, in particular the association between the pure quaterion portion and the relative expression profile.

Another type of quaternion model would map individual genes or pathways to the quaternion coefficients. For example, a cell-level gene set scoring method such as our Variance-adjusted Mahalanobois (VAM) [26] or Reconstruction Set Test (RESET) [27] method could be used to score three distinct gene sets (e.g., gene sets that capture different types of cytokine signaling, or gene sets that represent different cell type signatures) at each cell/location and these scores could then be used for coefficients of the vector part of the quaternion with the real part still representing sequencing depth. While this approach would only consider a subset of the expression profile, it would have the benefit of a more straightforward biological interpretation and a more comprehensive analysis could be realized via multiple quaternion mappings that are on different gene set combinations. In the remainder of this paper, we only explore the Algorithm 1 mapping and leave alternative representations to future work.

## 3 Quaternion model applications

This section details four different applications of the quaternion model detailed in Algorithm 1: visualization of cell state uncertainty, interpretation of cell state transitions, visualization of spatial transcriptomics (ST) data, and Fourier-based analysis. It is important to note that the first three applications do not strictly require a mapping to quaternions, i.e., they can be realized by the vector analysis of the projection of unnormalized count data onto uncentered PCs. The last application, however, can only be fully realized by a quaternion representation of the transcriptomic data. For this paper, we have just implemented the last two applications and leave the visualization of cell state uncertainty and characterization of cell state transtions to future work.

### 3.1 Visualization of cell state uncertainty

Single cell transcriptomics data is typically visualized by projecting normalized cells onto a 2D subspace with the first two PCs or first two UMAP dimensions common options. Although library size-based normalization does enable the comparative evaluation of cells/locations that may contain different amounts of mRNA or have different capture efficiencies/sequencing depths, it introduces an important limitation for standard visualization techniques (and other statistical analyses). Specifically, the uncertainty in the relative abundance values generated by normalization is inversely proportional to library size, however, this information is lost after normalization. Our approach for representing cells/locations as vectors in ℝ^3^ enables a visualization that captures both the data characteristics seen in a 2D projection of normalized data as well as uncertainty in the relative expression profile. Specifically, all cells/locations that share a similar relative expression profile will be distributed along a line that intersects the origin with the distance from the origin capturing relative library size. Cells/locations far from the origin therefore have a much more accurately inferred state than those close to the origin. This property can be used informally by users to guide the interpretation of the visualized scRNA-seq/ST data or formally to generate confidence regions for each cell/location according to a specific statistical model for the unnormalized counts.

### 3.2 Interpretation of cell state transitions

According to the quaternion model detailed in Section 2, the transition between cell states is represented by a rotation in ℝ^3^. As detailed in Section 1.2, this rotation can be represented by a rotation quaternion. An important benefit of this model is that the rotation axis defined by the rotation quaternion corresponds to a cell state and thus provides a biological interpretation for cell state transitions. More specifically, the rotation axis represents the cell state that is invariant to the associated transition, i.e., a cell in that state would be unaffected by whatever biological perturbation triggers a transition between those cell states. Although viewing cell state transitions in terms of an invariant or orthogonal state may not always be intuitive, it does provide a unique biological perspective on cell state differences.

An important application of this cell state transition model involves the intepretation of trajectories generated by pseudo-temporal ordering methods such as Monocle [20]. Specifically, the transitions between cells lying on a computed trajectory correspond to rotations and therefore have an interpretion via the associated invariant cell state. The quaternion rotation model also makes it possible to efficiently compute over the set of potential paths that connect cells in distal parts of the trajectory, e.g, cell at the root and cells at the leaves, which can be utilized to estimate the likelihood of the observed trajectory relative to the space of possible trajectories.

### 3.3 Visualization of spatial transcriptomics data

The standard 2D visualization of scRNA-seq data cannot be applied to ST data without losing spatial context. Visualization of ST data that preserves the spatial tissue structure is therefore usually performed for just a single gene at a time with color representing expression magnitude. Our proposed quaternion model enables a range of false color visualizations that capture both tissue architecture and a representation of entire transcriptional profile. Specifically, the four real coefficients associated with each quaternion, (*a, b, c, d*), can be mapped into red, green, blue (RGB) and *α* (i.e., opacity) values. One obvious mapping, which was employed by Sangwine [28] and Ell and Sangwine [8] to map from RGB values to quaternions sets the imaginary quaternion coefficients *b, c, d* to the red, green and blue values (see Ell and Sangwine for more details on the utility of this mapping for various image analysis tasks). We extend that mapping to also use the real quaternion coefficient *a* to represent the *α* opacity value. To realize this mapping, we rescale all quaternion coefficients to fall into a 0-to-1 scale, i.e., the minimum coefficient across all locations is set to 0 and the maximum to 1. This mapping has the benefit that the relative transcriptional profile of each ST location corresponds to a color with the opacity based on the library size. Specifically, locations with no reads will be fully transparent and those with the most reads will be fully opaque.

### 3.4 Spectral analysis using hypercomplex Fourier transformation

The genome-wide spectral analysis of scRNA-seq/ST data is challenging. For exploring the spectra of single cell transcriptomics in one dimension (e.g., pseudotime) or two dimensions (e.g., spatial location), a one or two dimensional discrete Fourier analysis is typically performed on the expression values for each gene separately. Alternatively, the projection of each cell onto a reduced dimension (e.g., PC or UMAP dimension) or pathway-based aggregation could be used but exploring the joint spectral characteristics of multiple gene expression variables is challenging. Ell and Sangwine [8] used a mapping from RGB color values to quaternions to support the application of a 2D quaternion-domain discrete Fourier transform [29] to color image data. As detailed by Ell and Sangwine, this quaternion-domain Fourier analysis can capture image features that are impossible to detect using separate Fourier transformations of each color component. Our proposed quaternion model enables the spectral analysis of scRNA-seq/ST data relative to a single variable (e.g., pseudo-time) or two variables (e.g., spatial location) to be performed on a genome-wide basis by used a 1 or 2D quaternion-domain discrete Fourier transformation applied to the quaternion representations of each cell/location. The output from such a discrete Fourier transform (especially the 2D transform of ST data) has a wide range of important applications [32], including spectral filtering, convolution-based analyses, matrix reconstruction [33], and registration of multiple matrices [34]. Examples of spectral filtering and convolution for ST data are explored in Section 4 below.

To realize this quaternion-domain Fourier transform using standard complex-domain discrete Fourier transform implementations, we can follow the approach of Ell and Sangwine [8]:

1. Rewrite the quaternions in the input vector **q** (for scRNA-seq data) or matrix **Q** (for ST data) in Cayley–Dickson form:

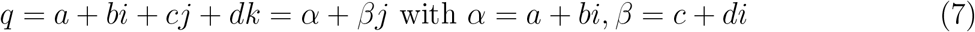
2. Create two new complex-values vectors (**c**_*α*_, **c**_*β*_) or matrices (**C**_*α*_, **C**_*β*_) that hold the *α* and *β* values from the Cayley-Dickson representation.
3. Compute the 1D or 2D complex-domain discrete Fourier transformations of **c**_*α*_, **c**_*β*_ or **C**_*α*_, **C**_*β*_. This can be realized using standard FFT implementations such as the *fft*() function in R. Let the output from these Fourier transformations be held in complex-valued vectors **f**_*α*_, **f**_*β*_ or matrices **F**_*α*_, **F**_*β*_.
4. Construct the quaternion-valued DFT output vector **f** or matrix **F** using the outputs of the complex-domain Fourier transformation, e.g., **f** = **f**_*α*_ + **f**_*β*_*j* or **F** = **F**_*α*_ + **F**_*β*_*j*.

## 4 Analysis of mouse brain spatial transcriptomics data

To illustrate the proposed quaternion model and the ST visualization and hypercomplex Fourier analysis applications, we analyzed a 10x Visium spatial transcriptomics dataset generated on mouse sagittal brain slices. This particular dataset is accessible as the *stxBrain* dataset in the *SeuratData* R package. Figure 2 shows the H&E stained section overlaid by a visualization of the Visium dots with color corresponding to the number of detected UMI counts.

**Figure 2:**
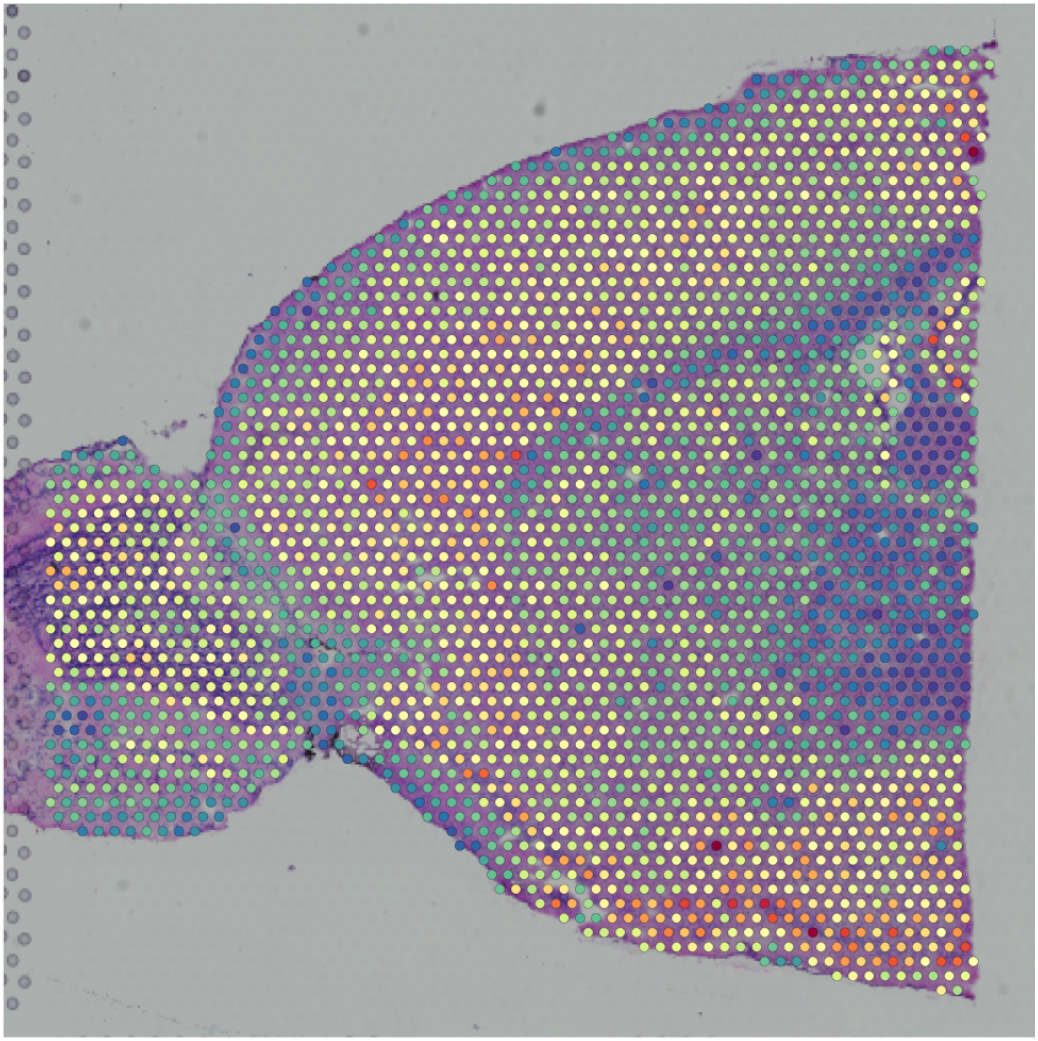
Visualization of 10x Visium spatial transcriptomics data for sagittal mouse brain slice. The H&E stained image is displayed behind a visualization of the Visium spots colored according to the number of UMI counts.

Processing of this ST data was realized using the Seurat framework [30] and QSC R package (available at https://hrfrost.host.dartmouth.edu/QSC). The QSC package implements the proposed quaternion mapping using the hypercomplex number functionality in the *onion* R package [31] and randomized SVD functionality in the *rsvd* R package [25]. Other important functions implemented by the QSC package include the quaternion-valued discrete Fourier transform (realized using the Cayley–Dickson approach detailed in Section 3.4), Fourier-based convolution of quaternion-valued matrices, generation and application of rotation quaternions, false color visualization of quaternion-valued matries (using the approach in Section 3.3), and mapping of ST data to quaternion matrices (see Section 4.1 below). The QSC package also includes a vignette that generates all of the mouse brain results included in this section as well as similar vignettes for a mouse kidney Visium ST dataset and a human ovarian cancer Vizgen MERSCOPE dataset.

### 4.1 Mapping of mouse brain ST data to quaternion matrix

To generate a quaternion matrix for the mouse brain ST data, the transcriptomic profile for each spot was first mapped to a quaternion using the approach defined by Algorithm 1 (note that this mapping ignores spot location). The quaternion values for each spot were then used to populate a 202-by-209 matrix whose rows represent evenly spaced vertical coordinates and whose columns represent evenly spaced horizontal coordinates on the Visium slide. For matrix elements not mapping to a spot, the quaternion value was initially set to (0, 0, 0, 0). Because this initial matrix representation was very sparse (only 6.4% of the elements mapped to a spot), a simple fill-in strategy was used that replaced empty elements with the average of adajent non-zero values. The resulting quaternion matrix representation of the mouse brain ST data is visualized in Figure 3 using the false color mapping detailed in Section 3.3.

**Figure 3:**
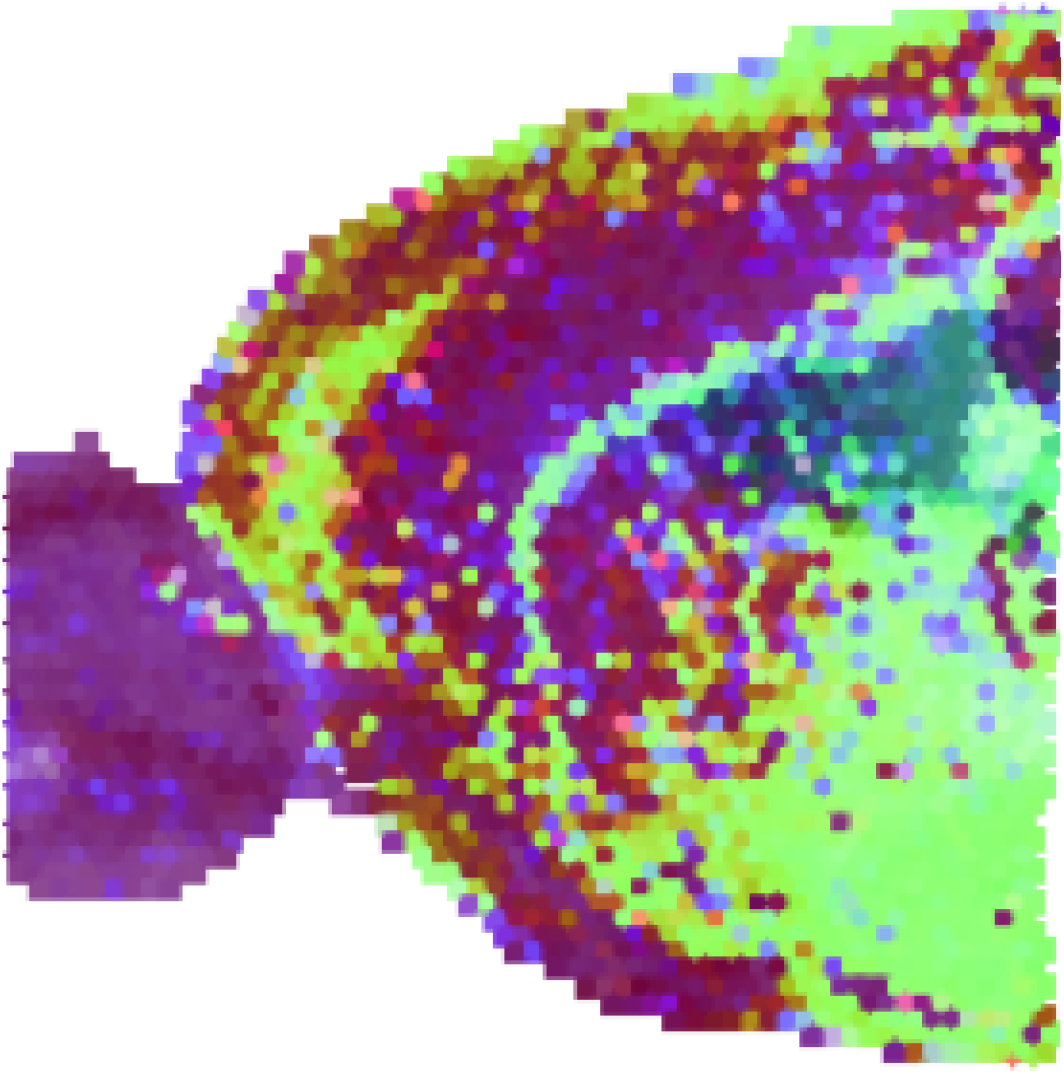
Visualization of quaternion mapping for sagittal mouse brain slice Visium ST data shown in 2.

### 4.2 Rotation of mouse brain quaternion matrix

An important benefit of the proposed quaternion model is that the vector part of each quaternion captures the relative expression profile of the associated cell/location and can be interpreted as a vector in ℛ^3^. As detailed in Section 1.2, the primary practical applications of quaternions is the efficient computation of 3D rotations. Figure 4 visualizes two different rotations of the quaternion representation of the mouse brain ST data. The left panel displays the result of rotating the vector components of each quaternion by 180 degrees about an axis defined by the *i* unit quaternion, i.e., a quaternion that has a coefficient of 1 for the *i* component and coefficients of 0 for all other components. Rotation of a vector quaternion about this axis keeps the *i* component constant and inverts the signs of the *j* and *k* components. In the context of our quaternion model, this will have the impact of keeping the projection onto the second non-centered PC constant and perturbing the second and third PC projections. The left panel visualizes the rotation of 180 degrees about an axis defined by the component of a quaternion for a spot near the choroid plexus. This type of rotation highlights regions of the tissue that are transcriptionally distinct from the choroid plexus.

**Figure 4:**
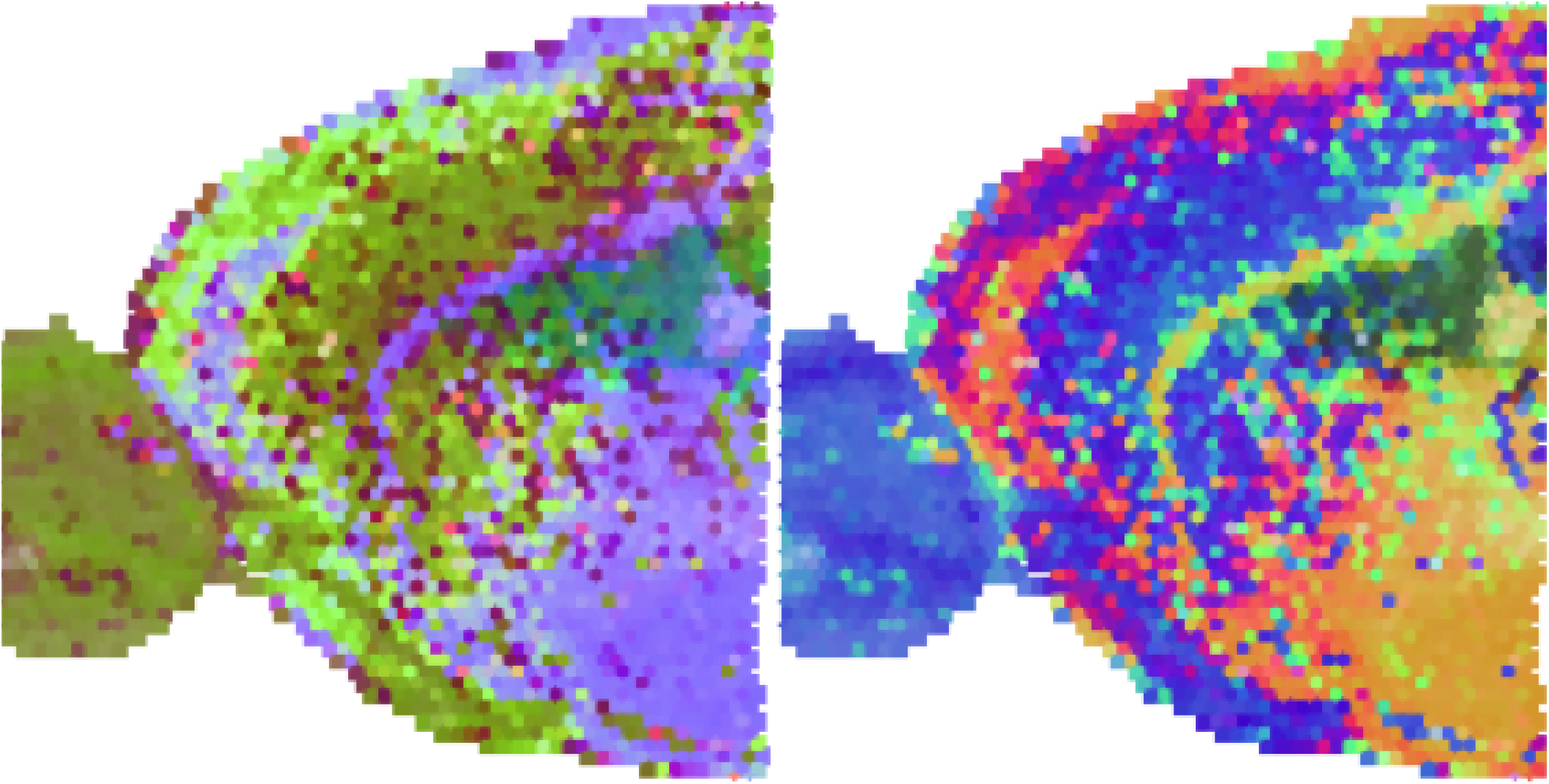
Visualization of the rotation of mouse brain quaternion matrix with about the quaternion *i* vector (left panel) or the quaternion associated with a spot near the choroid plexus.

### 4.3 Spectral filtering of mouse brain quaternion matrix

The proposed quaternion model enables the spectral analysis of ST data using a quaternion-domain 2D discrete Fourier transform. One type of simple Fourier-based analysis involves spectral filtering to remove specific frequency components. Specifically, the quaternion representation of the scRNA-seq or ST data is transformed into the frequency space using a quaternion-domain discrete Fourier transform (as detailed in Section 3.4), the quaternion values for undesired frequency components are set to 0 and result is transformed back using an inverse quaternion-domain discrete Fourier transform. The results generated by this type of spectral filtering of the mouse brain ST data are visualized in Figure 5 with the left panel displaying the output of low-pass filtering that removes the top 10 high frequency components and the right panel displaying the output of high-pass filtering that removes the bottom 10 low frequency components.

**Figure 5:**
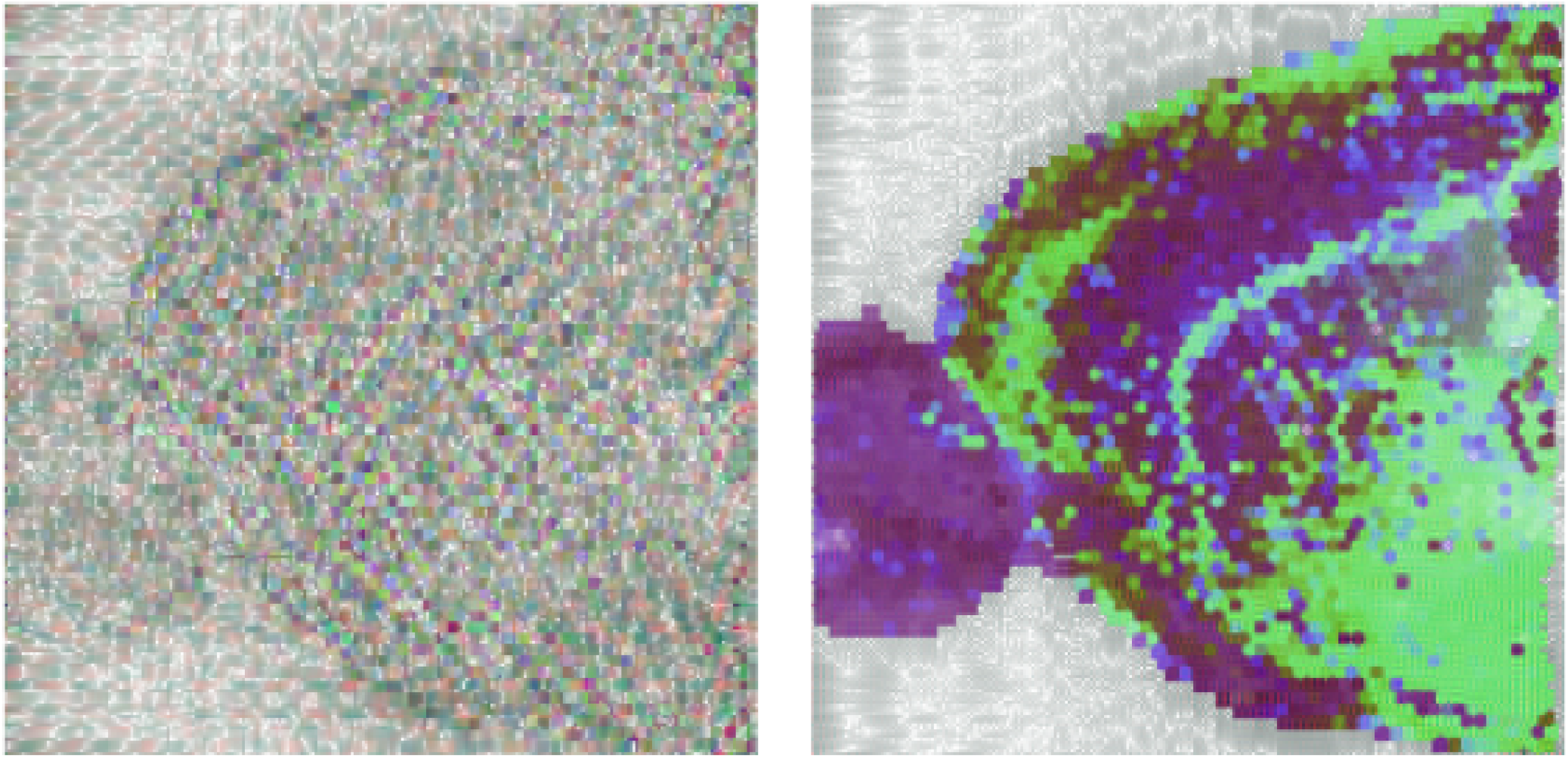
Visualization of the output of low-pass (left panel) or high-pass (right panel) filtering of the mouse brain quaternion matrix.

### 4.4 Convolution of mouse brain quaternion matrix with real-valued mask

The most important use case for quaternion-domain Fourier analysis is the computation of convolutions. Convolving image data with appropriate mask matrices is a critical step in many imaging processing pipelines and the QSC package provides support for the convolution of quaternion-valued matrices and generation of mask matrices that embed quaternion kernels. For the convolution of matrices generated according to the proposed quaternion model, the real component of the quaternions is ignored and the convolution is only applied to the vector portion that represents the relative transcriptomic profile.

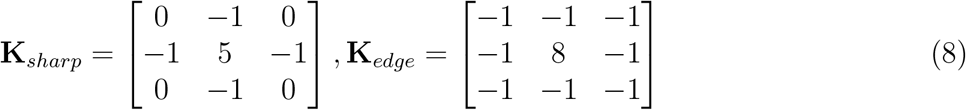

Figure 6 illustrates the output from the convolution of the quaternion matrix representation of the mouse brain ST data with a real-valued mask matrix. The left panel shows the output from convolution with a mask that embeds a 3-by-3 sharpening kernel (**K**_*sharp*_) and the right panel shows the output when the mask embeds a edge detection kernel (**K**_*edge*_; opacity is ignored to improve edge visualization).

**Figure 6:**
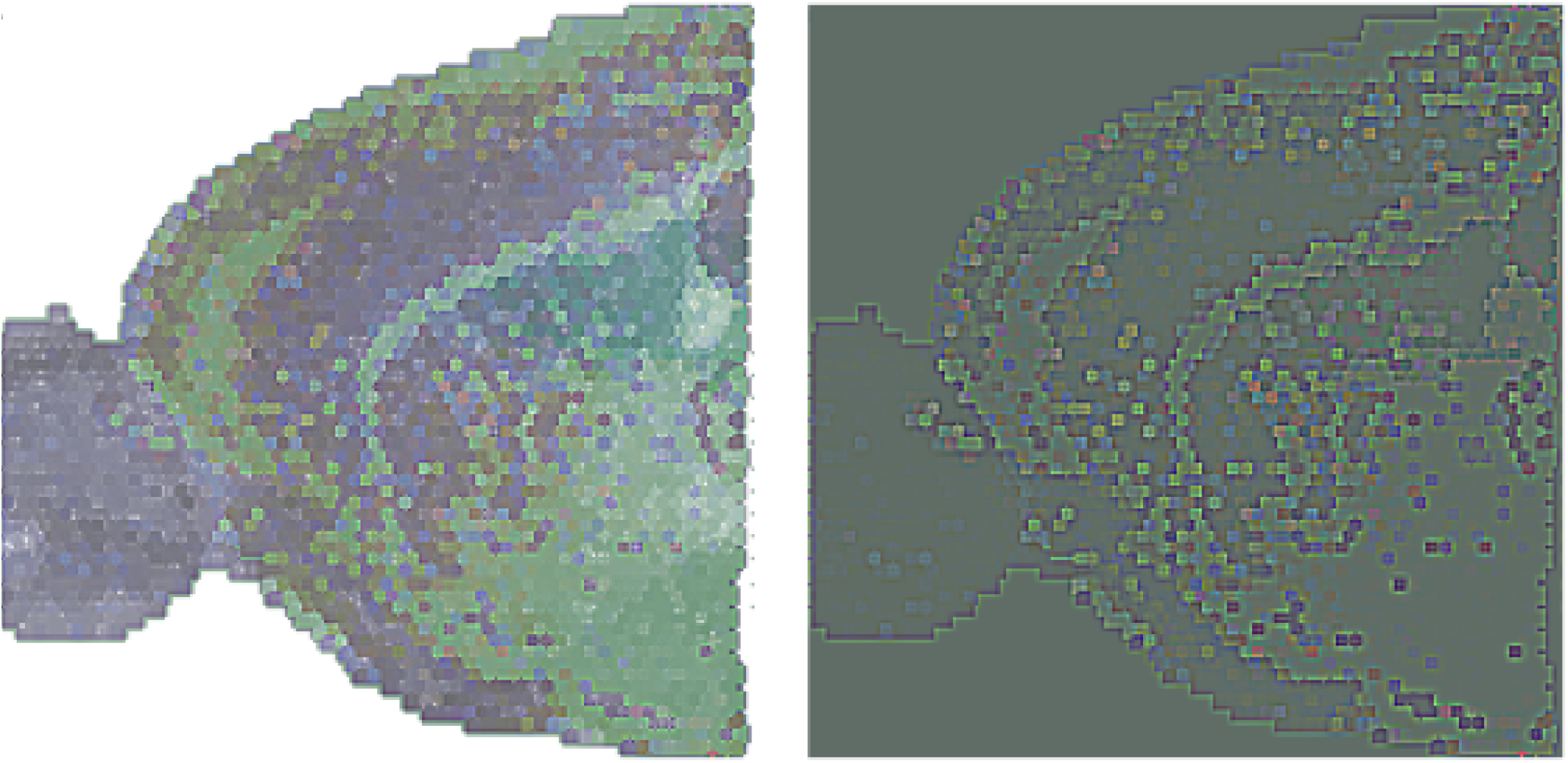
Visualization of the convolution of mouse brain quaternion matrix with masks that embed real valued sharpening (left panel) or edge detection (right panel) kernels.

### 4.5 Bi-convolution of mouse brain quaternion matrix with quaternion-valued masks

Although standard real-valued mask matrices that embed standard kernels (e.g., **K**_*sharp*_ and **K**_*edge*_) can be used, the true power of the quaternion model is realized by convolution of a quaternion-valued matrix representing an ST dataset with a quaternion-valued mask matrix. However, convolution with a quaternion-valued mask matrix encounters two complications:

- Quaternion multiplication is non-commutative so the left and right convolutions yield different results.
- Multiplication of two vector quaternions generates a quaternion with a real component.

Given these complications, design of a useful single quaternion convolution mask matrix is very challenging. These complications can be avoided through the bi-convolution of a matrix of vector quaternions (**X**) by left (**M**_*L*_) and right (**M**_*R*_) mask matrices:

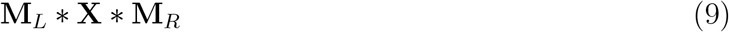

These type of structure has particular relevance in this context since rotation of a vector quaternion *q* is realized by pre and post-multiplication by a rotation quaterion *r* and its inverse via *rqr*^*−*1^. As explored by Ell and Sangwine [8], a version of edge detection can be performing via bi-convolution with mask matrices that embed the following 3-by-3 left and right quaternion-valued kernel matrices where *r* is a rotation quaternion:

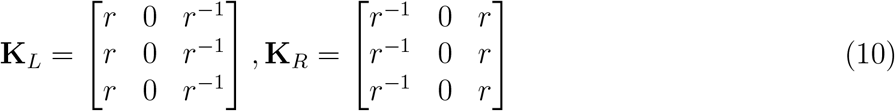

Bi-convolution with these kernels will perform edge detection in the space orthogonal to the rotation axis, i.e., changes in the direction of the rotation axis will be ignored and changes in directions orthogonal to the rotation axis will be highlighted. A straightforward example illustrated by the left panel of Figure 7 sets *r* to a 180 degree rotation about the *i* axis, which will detect transitions in the *j* and *k* components or projections of the spots onto the third and forth uncentered PCs. According to the false color visualization, these will be seen as edges in the green and blue portions of the spectrum. If the rotation axis is instead set *j*, which is visualized in the right panel of Figure 7, the convolution will identify edges in the *i* and *k* directions (or red and blue portions of the spectrum).

**Figure 7:**
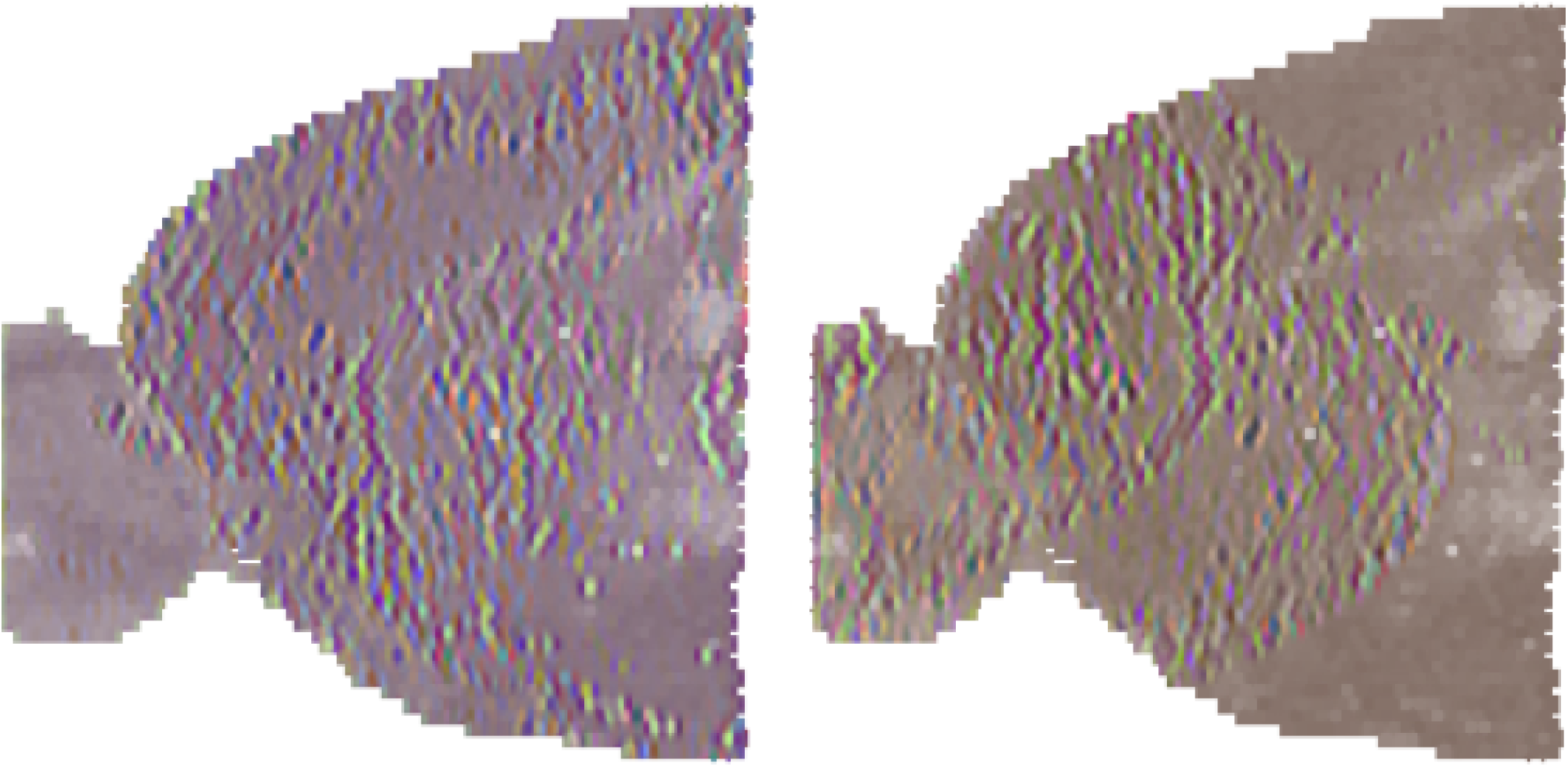
Visualization of the bi-convolution of mouse brain quaternion matrix with masks that embed quaternion-valued kernels that perform edge detection via rotation about a specific axis. For the left panel, the rotation axis is the *i* quaternion unit vector; for the right panel, the rotation axis is the *j* quaternion unit vector.

## 5 Limitations and future directions

There are three important limitations of our proposed quaternion model. First, the mapping from scRNA-seq/ST data to quaternions detailed in Algorithm 1 only uses the information in a rank 3 reduction of the unnormalized data, which will ignore a potentially substantial portion of the biological signal contained in the analyzed dataset. It is important to note in this context that the typical visualization/interpretation of scRNA-seq data involves a 2D projection, i.e., projection of the cells onto the first two UMAP dimensions, so significant practical utility can still be obtained from a quaternion model even if it misses biologically relevant characteristics of the data. Second, the non-commutative nature of quaternion multiplication makes the design and interpretation of analytical techniques such as convolution challenging. As illustrated through the ST example above, this challenge is mitigated by interpreting pure quaternions as vectors in ℛ^3^ and focusing on rotation-based operations. Finally, the false-color mapping outlined in Section 3.3 generates RGB mappings for a given quaternion that are relative to the range of quaternion values in the entire matrix. This relative mapping means that the color for a given element in the matrix may change after transformation of the matrix (e.g., convolution with another matrix) even when the quaternion associated with that element is unchanged.

In future work, we plan to address the first limitation by exploring alternative quaternion models (e.g., the use of multiple gene set-based models to analyze a single dataset) and the second limitation by designing quaternion-valued convolution masks that are not rotation-based and can be used without bi-convolution (e.g., masks that use the inverse of target quaternions in the kernel). For the third limitation, we will explore alternate RGB mapping approaches that maintain a consistent color for unchanged elements. We also plan to explore applications that were not detailed in this paper including the visualization of cell state uncertainty and the analysis of cell state transitions captured via pseudo-temporal ordering.

## 6 Conclusion

Quaternions are four dimensional hypercomplex numbers that have been extensively leveraged to represent three-dimensional rotations. In this paper, we detailed a novel approach for mapping single cell RNA-sequencing (scRNA-seq) and spatial transcriptomics (ST) data to quaternions. According to this model, the quaternion associated with each cell/location represents a vector in ℝ^3^ with vector length capturing sequencing depth and vector direction capturing the relative expression profile. This approach supports a number of potential applications including the visualization of cell state uncertainty, characterization of cell state transitions, visualization of ST data and, most importantly, the spectral analysis of scRNA-seq/ST data. To demonstrate the feasiblity of this approach, we mapped ST data for both Visium and Vizgen ST data to matrices of quaternions and explored a number of downstream analyses including the use of a quaternion-domain 2D discrete Fourier transform to compute convolutions and bi-convolutions of the ST quaternion matrix with real and quaternion-valued mask matrices. An R package supporting our proposed model and the hypercomplex Fourier analysis of ST data along with several example vignettes is available at https://hrfrost.host.dartmouth.edu/QSC.

## Acknowledgments

This work was funded by National Institutes of Health grants R35GM146586, R21CA253408, P20GM130454 and P30CA023108. We would like to acknowledge the supportive environment at the Geisel School of Medicine at Dartmouth where this research was performed.

